# Decoding isozyme-specific substrate recognition in protein arginine deiminases by proteome-wide citrullination mapping

**DOI:** 10.64898/2025.12.01.691579

**Authors:** Sophia Laposchan, Yi-Fang Yang, Kai-Han Chan, Rebecca Meelker Gonzalez, Wassim Gabriel, Mathias Wilhelm, Hui-Chih Hung, Chien-Yun Lee

## Abstract

Protein arginine deiminases (PADs) convert arginine to citrulline, altering protein structure and function. Of the five human isozymes, PAD1–4 are catalytically active with distinct tissue-specificities, yet isozyme-specific substrate recognition remains poorly defined. We profiled PAD1–4 substrate landscapes via mass spectrometry, identifying ∼30,000 citrullination sites across ∼5,500 proteins. Only 14% of sites were shared among all four, reflecting distinct sequence preferences: PAD1–2 showed broad specificity, whereas PAD3–4 favored arginines flanked by acidic or glycine residues. These preferences persisted over 10 min–16 h, indicating sequence context rather than temporal dynamics drives specificity. Mutation analysis of eleven PAD4 variants revealed Q346, G403, R639, and H640 as key determinants distinguishing substrate recognition from that of PAD2. This work provides the most comprehensive PADs substrate atlas to date, defining isozyme-specific motifs and molecular determinants, and guiding development of selective inhibitors and probes to interrogate citrullination mechanisms in health and disease.

## INTRODUCTION

Protein arginine deiminases (PADs) are a family of calcium-dependent enzymes that catalyse the post-translational conversion of peptidyl-arginine into citrulline, a process known as citrullination or deimination. This reaction removes the positive charge on the arginine side chain, thereby altering protein structure, dynamics, and interactions^1^. In humans, five PAD isozymes (PAD1–4, PAD6) are encoded by the PADI gene cluster; of these, only PAD1–4 show catalytic activity *in vitro*^1–3^.

The PAD family is highly conserved, sequence identity among human PADs ranges from ∼44% to ∼58%^4^ and key residues surrounding the substrate-binding pocket appear particularly well conserved across isozymes and species^5^. Such high homology suggests that subtle structural features, rather than gross fold differences, may underlie isozyme-specific substrate recognition. The isozymes exhibit distinct tissue distributions and subcellular localizations, suggesting distinct biological roles. PAD1 is predominantly expressed in the epidermis and uterus, PAD2 in the nervous and immune systems, and PAD3 in hair follicles. PAD4, uniquely among the active isozymes, contains a nuclear localization signal and is found both in the nucleus and cytoplasm, where it contributes to chromatin remodeling and gene regulation^1,6–8^. Dysregulated PAD activity is associated with various diseases such as rheumatoid arthritis (RA), multiple sclerosis, Alzheimer’s disease, and a variety of cancers^9–12^, rendering the PAD family important both for basic biology and therapeutic intervention.

Despite this broad interest and the known biological roles of PADs, the molecular basis of how individual isozymes select their substrates remains incompletely defined. Early biochemical studies used simple arginine derivatives or short synthetic peptides to measure activity (e.g., Nomura 1992^13^) but lacked the broader sequence and structural context of full-length proteins^14–19^. Subsequent work assessed in-cell or in-tissue citrullination patterns (e.g., in neutrophils or RA synovium) but these often involve mixed isozyme inputs^20,21^, limited site resolution^21^, or are constrained to a subset of PADs^20,22,23^. For example, studies showed that PAD2, PAD3 and PAD4 have distinct specificities against intracellular actin or histone H3^21^, arguing for isozyme-intrinsic selectivity beyond cellular localisation. Likewise, more recent analyses attempted substrate profiling of PAD2 and PAD4 using artificial peptide libraries or combinatorial approaches, yet these remain limited in scope^24^. Collectively, these efforts highlight key knowledge gaps: (i) no systematic, proteome-wide comparison of all catalytically active human PADs under uniform conditions; (ii) limited understanding of how binding-pocket homology translates into divergent substrate recognition; (iii) a lack of residue-level substrate mapping across a broad protein repertoire; and (iv) insufficient linkage between kinetic/ peptide-based assays and complex proteome-scale citrullination outcomes.

Technical hurdles have also hampered progress in the field. Most conventional detection methods (e.g., immunoblotting or antibody-based enrichment) act at the protein level and cannot localize citrullination sites. Even mass spectrometry-based bottom-up proteomics, which offers site-level resolution, faces challenges for citrullination: the mass shift (+0.984 Da) is identical to that of asparagine/glutamine deamidation, making false positives a major concern^25,26^. Recent advances in enrichment strategies^27–29^, computational analysis^30–32^ have improved confidence in detecting citrullination sites, but their application across PAD isozymes remains sparse.

Here, we provide a systematic, isozyme-wide comparison of human PAD1–4 by combining controlled *in vitro* incubation of human cell lysates with recombinant enzymes, deep-coverage LC–MS/MS and advanced computational validation. Importantly, by incubating lysates under uniform, saturating enzyme and calcium conditions we aimed to isolate intrinsic substrate recognition potential from confounding factors such as isozyme expression levels, localization or intracellular calcium concentrations. We further leveraged deep-learning-based rescoring to elevate site-level confidence^32^. With this approach we mapped ∼30,000 citrullination sites across ∼5,500 of proteins, enabling direct comparison of substrate repertoire, sequence motif, and structural context among PAD isozymes, setting the stage for mechanistic dissection of how binding-pocket homology and peripheral residues dictate specificity. To connect global substrate patterns to underlying molecular determinants, we complemented the proteome-wide mapping with a systematic analysis of eleven PAD4 mutants, enabling identification of key residues that drive isozyme-specific substrate recognition.

## RESULTS

### Proteome-Wide Citrullination Mapping Delineates PAD Isozyme-Specific Substrate Landscape

Enzyme kinetic assays provide valuable parameters for substrate binding and catalytic efficiency using single substrates of PAD, but they rarely capture isozyme-specific substrate selection across a complex proteome^18^. To address this, we performed a proteomic workflow to globally map PAD-catalyzed citrullination sites from human proteome (**Fig. 1A**). Briefly, cell lysates from two cell lines, HeLa and H4, were prepared under mild conditions to preserve protein structure and incubated with recombinant PAD1–4. After digestion and basic reverse-phase tip fractionation, peptides were analyzed by LC-MS/MS. Database search results were processed with Prosit-Cit, a machine-learning rescoring model predicting MS/MS spectra of citrullinated peptides to enhance precision of the identification^32^.

**Figure 1.**
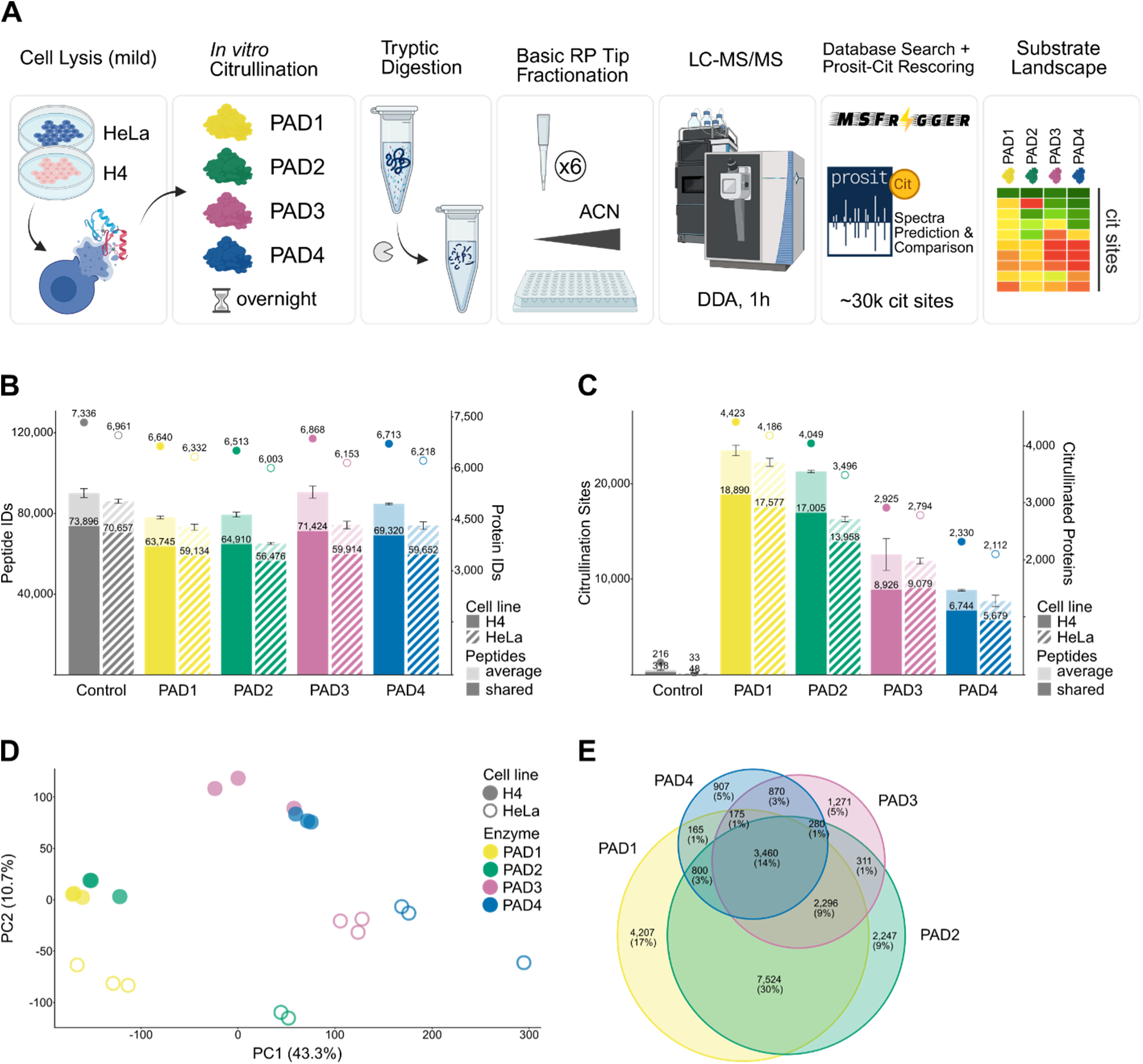
Proteome-wide citrullination mapping delineates PAD isozyme-specific substrate landscape. (A) Schematic overview of the workflow. (B) Total number of peptides and proteins identified per experiment. Light bars represent the average number of peptides across replicates (n = 3 for all experiments except HeLa PAD2, where n = 2), with standard deviations shown as error bars. Dark bars indicate peptides shared across all replicates, and dots represent shared proteins. (C) Number of citrullinated sites and proteins identified for each PAD isozyme compared to control (no enzyme). (D) Principal component analysis (PCA) of all citrullination sites identified across the dataset. (E) Overlap of citrullination sites catalyzed by PAD isozymes in H4 cell lysate. Sites shared exclusively between PAD1 and PAD3 (263 sites) or PAD2 and PAD4 (87 sites) were omitted to maintain proportional visualization.

This approach identified ∼85,000 peptides from ∼7,300 proteins in H4 and ∼75,000 peptides from ∼6,900 proteins in HeLa, with high reproducibility across replicates (**Fig. 1B**). In total, 29,485 citrullination sites across 5,480 proteins were detected (**Fig. 1C; Sup. Table 1**). Given the *in vitro* nature of our workflow, the observed citrullinome likely exceeds the physiological set of citrullinations reported previously^20,22,26^, as not all sites are expected to occur *in vivo*. Nevertheless, the breadth of substrates in whole-cell lysates provides a powerful framework to assess isozyme specificity under competitive conditions, generating the most comprehensive PAD substrate resource to date. Within this resource, PAD1 produced the most extensive citrullination (18,890 sites in H4, 17,577 sites in HeLa), followed by PAD2, whereas PAD3 and PAD4 modified substantially fewer sites (∼9,000 and ∼6,000 sites, respectively). At the protein level, PAD1 targeted over 4,400 proteins in H4 lysate compared to ∼2,300 for PAD4. These endpoint reactions indicate intrinsic differences in substrate promiscuity: PAD1 and 2 act broadly, while PAD4 is the most selective.

**Table 1.** Enzymatic activity of PAD2, PAD4, and PAD4 mutants towards model substrate BAEE. *K*_m_ = Michaelis constant. *k*_cat_ = turnover number. Values provided are mean ± standard deviation across triplicate results.

| | $K_m, \text{BAEE}$ (mM) | $k_{cat}$ ($\text{s}^{-1}$) | $k_{cat}/K_m, \text{BAEE}$ ($\text{mM}^{-1}\text{s}^{-1}$) |
| --- | --- | --- | --- |
| PAD2 WT | $0.12 \pm 0.01$ | $0.95 \pm 0.03$ | $7.92 \pm 3.00$ |
| PAD4 WT | $0.66 \pm 0.05$ | $5.98 \pm 0.14$ | $9.06 \pm 2.80$ |
| Mutant A (Q346R) | $0.93 \pm 0.07$ | $6.09 \pm 0.15$ | $6.55 \pm 2.14$ |
| Mutant B (R374G) | $0.26 \pm 0.01$ | $5.15 \pm 0.07$ | $19.81 \pm 7.00$ |
| Mutant C (G403S) | $0.32 \pm 0.08$ | $7.10 \pm 0.45$ | $22.19 \pm 5.63$ |
| Mutant D (R639F) | $0.17 \pm 0.03$ | $6.12 \pm 0.20$ | $36.00 \pm 6.67$ |
| Mutant E (H640L) | $0.30 \pm 0.04$ | $6.21 \pm 0.23$ | $20.70 \pm 5.75$ |
| Mutant AD | $0.12 \pm 0.04$ | $3.25 \pm 0.18$ | $27.08 \pm 4.50$ |
| Mutant CE | $0.16 \pm 0.02$ | $6.56 \pm 0.12$ | $41.00 \pm 6.00$ |
| Mutant DE | $0.06 \pm 0.04$ | $2.39 \pm 0.18$ | $39.83 \pm 4.50$ |
| Mutant AC | $0.33 \pm 0.06$ | $3.20 \pm 0.14$ | $9.70 \pm 2.33$ |
| Mutant ADE | $0.04 \pm 0.01$ | $2.40 \pm 0.02$ | $60.00 \pm 2.00$ |
| Mutant ACDE | $0.03 \pm 0.00$ | $2.94 \pm 0.03$ | $98.00 \pm 0.00$ |

Citrullinated peptides spanned the full intensity range of identified peptides, closely following the global distribution (**Sup Fig. 1A,B**), confirming our workflow enables detection over a wide dynamic range across low- and high-abundance peptides. PAD3/4 substrates tended to fall into lower-intensity bins, consistent with narrower and potentially lower-abundance substrate pools.

Due to the setting of our workflow, multiple modification on the same protein was frequent, with over 70% of proteins harboring more than two citrullination sites (**Sup Fig. 1C,D**). PAD1/2 showed the greatest propensity for multi-site events, suggesting relaxed sequence or structural requirements. For particularly large proteins such as plectin (∼4,700 aa, 175 sites), epiplakin

(∼5,000 aa, 137 sites), and dynein heavy chain (∼4,700 aa, 116 sites) extensive modification was observed. Well-established targets including vimentin (21 sites), α-enolase (12 sites), and histone H3.2 (7 sites) were also citrullinated at multiple residues, including previously undescribed sites (**Sup. Table 1**).

To compare substrate landscape across isozymes and cell lines, we performed principal component analysis. PAD1 and PAD2 clustered closely, distinct from PAD3 and PAD4 (**Fig. 1D**). These enzyme-driven differences exceeded the variation between HeLa and H4 proteomes, indicating that substrate selectivity is largely driven by intrinsic isozyme properties rather than proteome background. Hierarchical clustering confirmed two major groups: PAD1/2 versus PAD3/4 across both cell lines, with the exception of HeLa PAD2 samples that grouped closer to PAD3/4 (**Sup Fig. 1E**).

Direct comparison of modification sites captured this partitioning (**Fig. 1E; Sup. Fig. 1F,G**). PAD1 and PAD2 shared 56% of all sites (14,080 sites in H4 lysate), with ∼30% (7,524 sites) unique to this pair. PAD3 and PAD4 showed a minimal overlap exclusive to them (∼3%, 870 sites). Only ∼14% (3,460 sites) of citrullination sites were shared across all isozymes, representing a modest core citrullinome. These results align with the observed clustering and demonstrate that PAD isozymes operate with largely non-redundant substrate repertoires.

### Motif and Structural Analyses Reveal Distinct Preferences of PAD1/2 versus PAD3/4

To explore the basis of isozyme-specific citrullination, we examined the sequence and structural features of modified sites, asking whether differences in local motifs or structural context contribute to the distinct substrate repertoires. Motif enrichment analysis was used to compare amino acid frequencies within a ±5 residue window around citrullinated arginines to those surrounding all arginines in the respective proteome^33^ (**Fig. 2A; Sup. Table 1**). PAD1 and PAD2 showed broad enrichment for both basic (K) and acidic (D, E) residues across the flanking region, with additional preference for hydrophobic residues such as I, L, V, and A. In contrast, PAD3 and PAD4 displayed more focused motifs: a strong bias for D, S, or T at the –1 position and E, D, G, or N at +1. While distal positions also showed enrichment for D, E, and K, the narrow composition adjacent to the modified arginine suggests stricter substrate requirements for PAD3/4. Similar trends were observed in both HeLa and H4 (**Fig. 2A; Sup Fig. 2A**), with subsequent analyses focused on H4 lysate.

**Figure 2.**
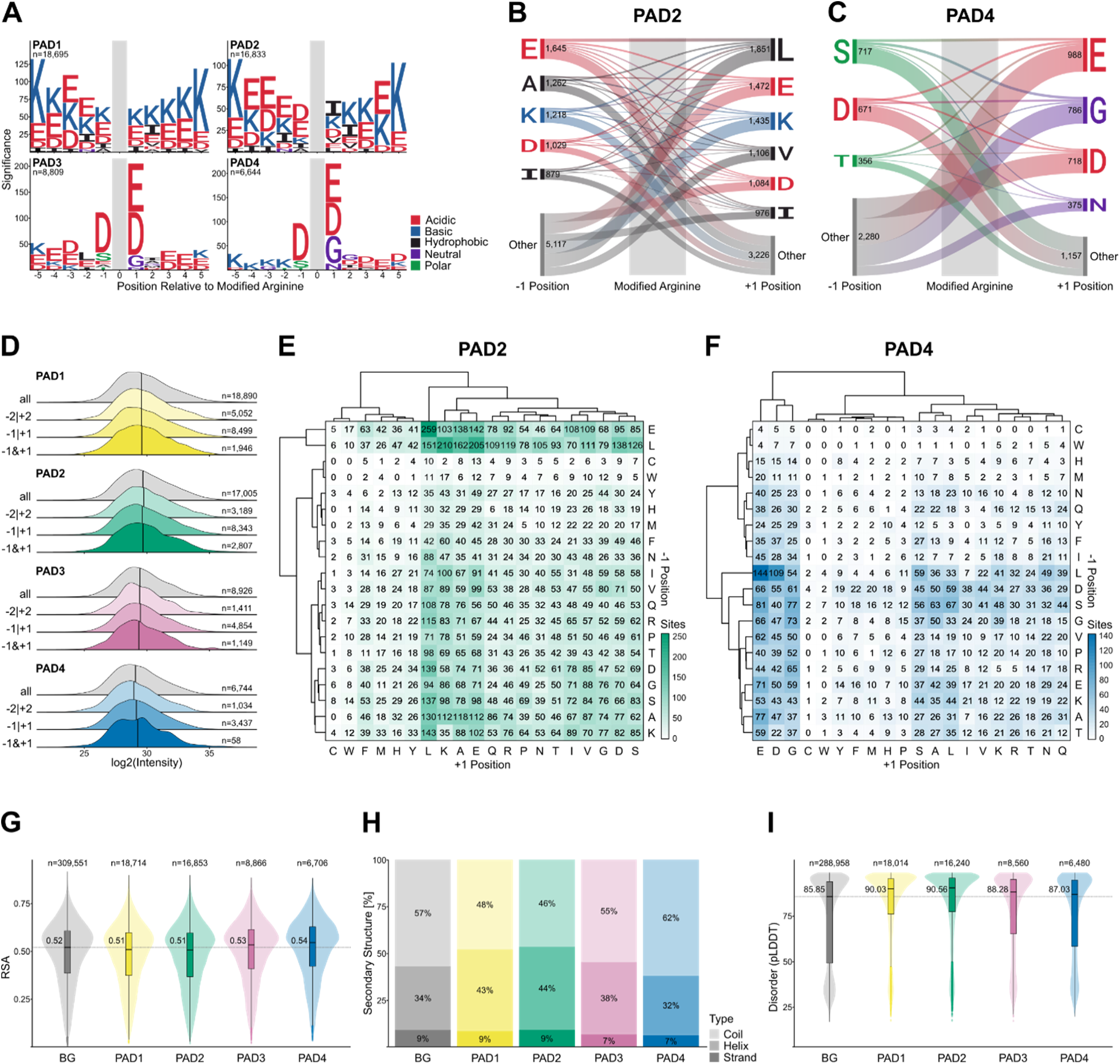
Motif and structural analyses reveal distinct preferences of PAD1/2 versus PAD3/4. (A) Motif enrichment analysis of citrullination sites generated by each PAD isozyme in H4 lysate compared to background of all identified proteins. (B, C) Co-occurrence of residues flanking the modified arginine for PAD2 and PAD4. Only sites containing a favored residue at either position, as defined by motif analysis in (A), are included. Connections indicate the frequency of residue pairings between positions −1 and +1. (D) Intensity distributions of all citrullination sites compared to sites containing favored residues at −2 or +2 (−2|+2), −1 or +1 (−1|+1), or both −1 and +1 (−1&+1) positions. (E, F) Frequency of amino acid combinations at −1 and +1 positions in PAD2- and PAD4-modified sites. Rows and columns are hierarchically clustered by Euclidean distance. (G) Predicted relative solvent accessibility (RSA) of modified arginine residues compared to all arginines within identified proteins (background, BG). (H) Ratio of arginine residues within coil, helix, and strand regions for PAD-modified versus all arginines (BG). (I) Distribution of AlphaFold pLDDT values for PAD-modified arginines compared to all arginines within identified proteins (BG).

We next asked whether PADs require combinations of preferred residues flanking the target arginine. Most citrullination sites contained only a single favorable residue on either side, without a consistent requirement for paired residues (**Fig. 2B,C; Sup. Fig. 2B,C**). This suggests that a single stabilizing interaction at one adjacent position is often sufficient for catalysis. Consistent with this, intensity distributions of peptides containing preferred residues at positions -2 or +2, - 1 or +1, or both -1 and +1 did not differ significantly (**Fig. 2D**).

Analysis of residue pair co-occurrence further highlighted isozyme differences. PAD1/2 tolerated a wide range of residue combinations, such as E or L at –1 paired with L, K, A, or E at +1 (**Fig. 2E; Sup. Fig. 2D**), reflecting mixed usage of charged and hydrophobic environments. PAD3/4, by contrast, showed a much narrower motif landscape, with only a few combinations (e.g., L at –1 with E or D at +1) appearing repeatedly (**Fig. 2F; Sup Fig. 2E**). This restricted motif repertoire reinforces the more selective substrate profile of PAD4.

We then examined structural features of proteins bearing citrullination sites using NetSurfP-3.0^34^ and AlphaFold-predicted structures^35^, to explore the structural context of their localization. PAD3/4 sites were more often located in solvent-accessible regions compared to PAD1/2, though the difference was modest (**Fig. 2G**). Secondary structure predictions indicated that PAD1/2 more frequently targeted arginines within helices, whereas PAD3/4 preferentially modified residues in coil regions (**Fig. 2H**). This was further supported by AlphaFold pLDDT scores: PAD3/4-modified sites had lower confidence values, consistent with localization in disordered regions, while PAD1/2 sites were enriched in higher-confidence, ordered domains (**Fig. 2I**). Together, these data indicate that PAD1/2 tolerate broader sequence and structural contexts, whereas PAD3/4 preferentially act on accessible residues within disordered regions.

### Time-Resolved Profiling Demonstrates Consistent Substrate Specificity of PAD2 and PAD4

Overnight incubations capture the full substrate range *in vitro*, but PADs may act transiently *in vivo*. To examine whether substrate preferences shift over time, we focused on PAD2 and PAD4, which represent the two major substrate-selectivity subtypes: PAD2 as broadly reactive and PAD4 as the most selective. H4 lysates were incubated with each isozyme at five time points (10 min to 16 h) to compare early versus late substrate engagement. Both enzymes showed increasing numbers of modified sites over time (**Fig. 3A; Sup. Table 2**). PAD2 exhibited rapid kinetics, generating more than half of its endpoint sites within 10 minutes and plateauing by 3 hours. PAD4 accumulated sites more gradually, with ∼2,000 additional modifications (∼22% of endpoint total) detected between 3 h and overnight, consistent with continued access to selective targets.

**Figure 3.**
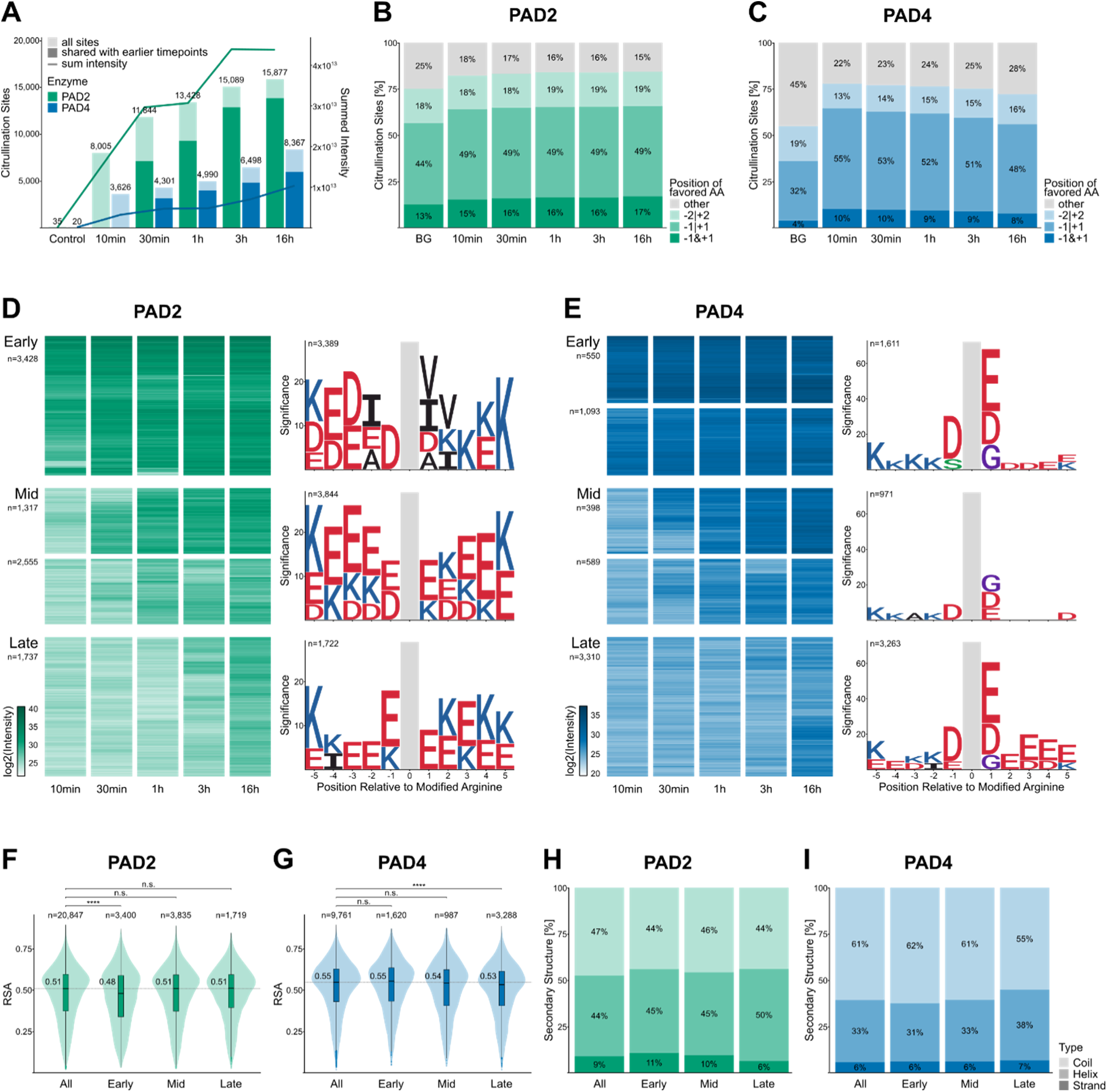
Time-resolved profiling demonstrates consistent substrate specificity of PAD2 and PAD4. (A) Number of identified citrullination sites (bars) and their summed intensity (lines) across incubation times with PAD2 and PAD4. Light bars denote total sites per timepoint; dark bars indicate sites shared with previous timepoints. Control sample was incubated without enzyme. (B, C) Proportion of citrullination sites containing favored residues (as defined in Fig. 2A) at −2/+2, −1/+1, or both −1 and +1 positions for PAD2 and PAD4. Background (BG) represents the proportion of arginine residues bearing favored neighboring residues (regardless of modification) within all proteins identified in H4 lysate. AA, amino acid. (D, E) Hierarchical clustering (Euclidean distance) of citrullination sites reveals early, intermediate, and late conversion clusters for PAD2 and PAD4, respectively. Right panels show motif enrichment for sites within each cluster. (F, G) Predicted relative solvent accessibility (RSA) of PAD2- and PAD4-modified arginines across all timepoints (All) and within Early, Mid, and Late clusters defined in (D)/(E). Statistical significance was assessed using the Wilcoxon test (**** = p ≤ 0.0001; n.s., not significant). (H, I) Ratio of PAD2-and PAD4-modified arginines predicted to reside in coil, helix, or strand regions across all timepoints (All) and within Early, Mid, and Late clusters as defined in (D)/(E).

To assess whether sequence preferences changed over time, we first compared the proportion of citrullination sites with favored flanking residues (–2 to +2) identified in the endpoint motif analysis. PAD2 showed stable distributions, with ∼15–17% of sites carrying favored residues on both sides of the citrullinated arginine, ∼49% on one side, and ∼18–19% at distal –2 or +2 positions (**Fig. 3B, C**). PAD4 displayed a subtle decline in sites containing favored residues at –1 and/or +1 over time, suggesting gradual depletion of optimal motifs. Clustering sites by temporal intensity profiles further highlighted these differences (**Fig. 3D, E**). PAD2 initially favored hydrophobic residues at –2, +1, and +2, but later clusters shifted toward acidic and basic residues (D, E, K), consistent with its broad endpoint motif (**Fig. 3D; Sup Fig. 3A**). PAD4, by contrast, maintained its canonical –1/+1 signature across early, mid, and late clusters and sites identified across all timepoints (**Fig. 3E; Sup Fig. 3B**).

We next examined whether structural context shifted with time. For PAD2, early-modified sites had slightly lower solvent accessibility than later sites (**Fig. 3F**), suggesting an early preference for less exposed but hydrophobic residues, potentially stabilized by local interactions. PAD4 showed the opposite trend: later-modified sites had lower accessibility (**Fig. 3G**), consistent with progressive engagement of less accessible but motif-compatible targets after the most accessible sites were consumed.

Secondary structure predictions showed that PAD2 substrates remained consistently distributed across helices, strands, and coils throughout the time course (**Fig. 3H**). PAD4, however, increasingly citrullinated residues in helices and strands at later points, whereas early-modified sites were more frequently located in coils (**Fig. 3I**). This indicates that PAD4 progressively engages structurally constrained substrates, but only when they conform to its strict sequence requirements. Together, these analyses highlight that PAD2 rapidly modifies a broad range of targets and shows a mild early bias for hydrophobic environments, whereas PAD4 maintains strict sequence selectivity and gradually expands to less accessible, more structured substrates over time.

### Single Point Mutations on PAD4 Expand Substrate Repertoire

Given PAD4’s narrow substrate profile compared to PAD2’s broader specificity, we hypothesized that residues near the substrate-binding pocket provoke these differences. Based on prior structural studies (Q346 in Arita *et al.*^36^; R374 and R639 in Lee *et al.*^37^) and sequence divergence between PAD isozymes (**Fig. 4A**), we selected five candidate residues (Q346, R374, G403, R639, H640) located within 5 Å of the active site or in close proximity according to linear sequence (**Fig. 4B**). Each was substituted with the PAD2 counterpart to generate five single-point PAD4 mutants: A (Q346R), B (R374G), C (G403S), D (R639F), and E (H640L). These variants were tested using the same proteomics workflow as described (**Fig 1A**), alongside kinetic assays with the model substrate BAEE (Benzoyl-L-arginine ethyl ester) to measure binding and catalytic efficiency.

**Figure 4.**
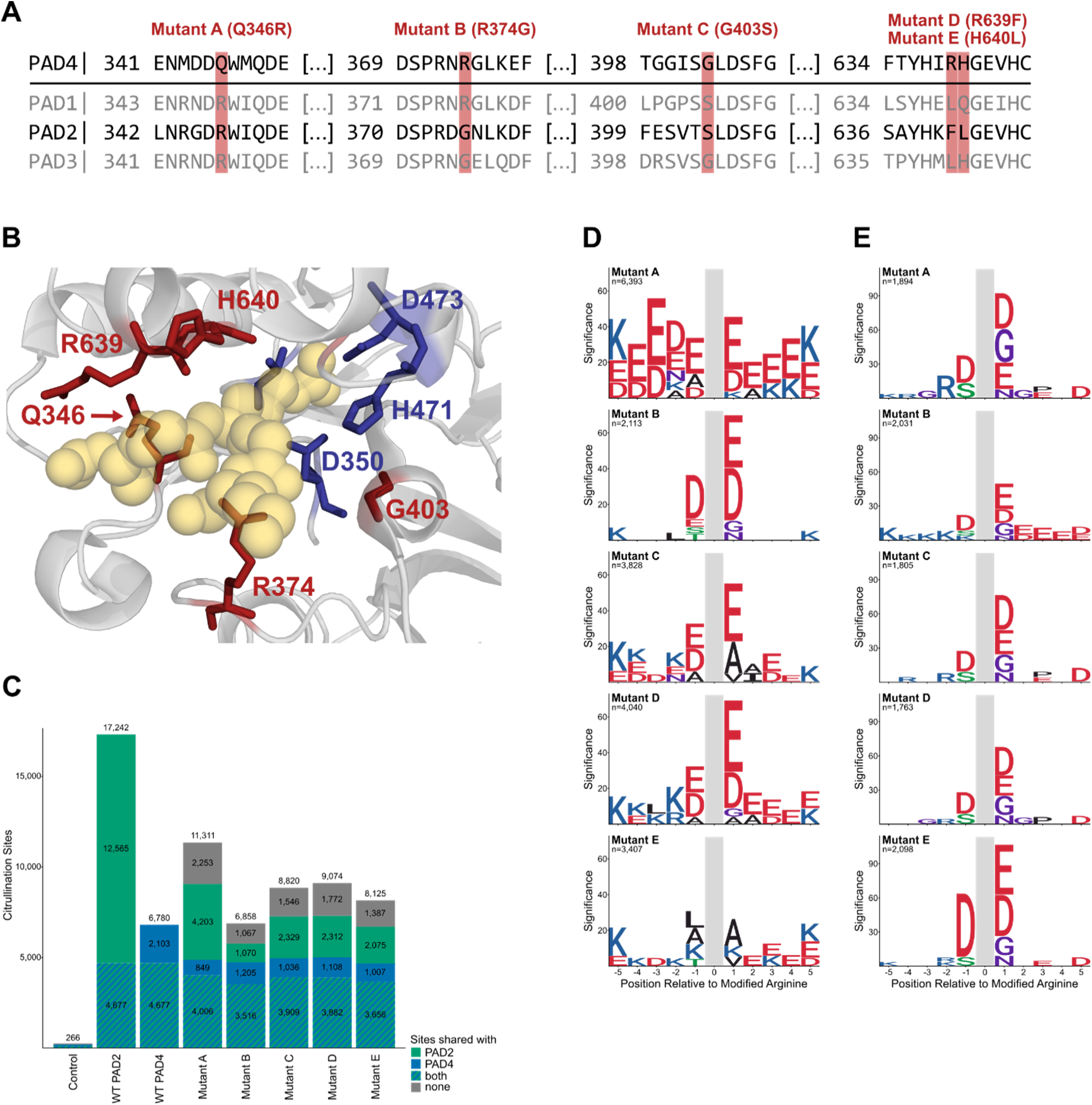
Single point mutations on PAD4 expand substrate repertoire. (A) Sequence alignment excerpts of PAD1–4 near known substrate-binding regions. Residues selected for mutagenesis are highlighted in red. (B) Crystal structure of PAD4 (grey) in complex with histone H4 substrate (yellow) (PDB ID: 2DEY). Catalytic residues are shown in blue, and mutated residues in red. (C) Number of citrullination sites identified for each enzyme or PAD4 mutant compared to control (no enzyme). Shading indicates sites shared with wild-type PAD2, PAD4, or both. (D) Motif enrichment analysis of sites newly generated by each mutant (i.e., not shared with wild-type PAD4). (E) Motif enrichment analysis of sites lost in each mutant (i.e., present in wild-type PAD4 but absent in the corresponding mutant).

All single-point mutants retained enzymatic activity toward BAEE (**Table 1**), and displayed either comparable or enhanced activity compared to wild-type (WT) PAD4. While Mutant A exhibited slightly reduced substrate affinity, Mutants B and E showed roughly double the catalytic efficiency of WT PAD4 (19.81 and 20.70 mM⁻¹s⁻¹, respectively, compared to 9.06 mM⁻¹s⁻¹), and Mutant D demonstrated the most significant improvement, almost 4-fold greater efficiency (36 mM⁻¹s⁻¹). These results indicate that alterations at select sites can markedly affect catalytic performance toward simple substrates.

However, proteome-wide assays revealed a more complex outcome (**Fig. 4C; Sup. Table 3**). All mutants except Mutant B citrullinated substantially more sites than WT PAD4. Mutant A modified 11,311 sites, representing a ∼66% increase over WT. Mutants C, D, and E also generated ∼20–30% more sites than WT PAD4. By contrast, Mutant B, highly efficient toward BAEE, produced only a marginal increase in site count. These findings highlight that catalytic performance on a single substrate does not reliably predict enzyme behavior across thousands of competing proteins in a proteome. Comparison with WT PAD2 revealed that many of the additional sites introduced by PAD4 mutants overlapped with PAD2-specific targets absent from WT PAD4. Between 2,000 and 4,000 “gain” sites matched WT PAD2 but not PAD4, whereas the number of sites shared with both PAD2 and PAD4 (∼3,500–4,000) or unique to WT PAD4 (∼1,000) remained constant across all mutants. Thus, individual substitutions enabled PAD4 to expand its repertoire into the PAD2 target space without eliminating its native profile.

Motif analyses further supported this shift. Across all sites, the canonical PAD4 motif (D at –1; E, D, G at +1) remained dominant (**Sup. Fig. 4**). However, gain sites unique to mutants revealed relaxed preferences (**Fig. 4D**). Mutant A accepted broader enrichment for acidic and basic residues (E, D, K) around the target arginine, resembling PAD2’s profile, while Mutant C and E incorporated hydrophobic neighbors such as A, L, I, and V. Conversely, sites lost in mutants but present in WT PAD4 consistently displayed the strict canonical motif (**Fig. 4E**). Together, these results show that single substitutions loosen PAD4’s sequence requirements, allowing access to additional PAD2-like sites while maintaining its core specificity.

### Multiple PAD4 Mutations Drive Broad Substrate Recognition Similar to PAD2

While single mutations showed modest shifts in substrate motifs, particularly at Q346 (site A), G403 (site C), R639 (site D), and H640 (site E), we hypothesized that these sites act collectively to govern PAD4 substrate recognition. To test this, we engineered multiple mutants incorporating combinations of these residues. This included four double mutants, AC (Q346R|G403S), AD (Q346R|R639F), CE (G403S|H640L), and DE (R639F|H640L), a triple mutant ADE (Q346R|R639F|H640L), and a quadruple mutant ACDE (Q346R|G403S|R639F|H640L). All variants retained catalytic activity toward the model substrate BAEE, with most showing improved affinity and several reaching or exceeding the binding affinity of WT PAD2 (**Table 1**). Although *k*_cat_ values generally remained lower than WT PAD4, enhanced substrate affinity drove markedly increased catalytic efficiencies. The triple mutant ADE achieved an 6.6-fold improvement (60 mM⁻¹s⁻¹), and the quadruple mutant ACDE exceeded a 10.8-fold gain (98 mM⁻¹s⁻¹ vs. 9.06 mM⁻¹s⁻¹ for WT PAD4).

In proteome-wide assays, however, kinetic gains did not always correlate with expanded substrate profiles (**Fig. 5A; Sup. Table 3**). Double mutant CE for instance produced only 7,438 sites, slightly fewer than either single mutant C (8,820) or E (8,125), suggesting context-dependent effects. Mutant DE similarly showed no additive increase compared to Mutant D alone. By contrast, inclusion of Q346R (site A) consistently drove substantial expansion. Double mutants AC and AD generated 9,176 and 9,530 sites, respectively, slightly fewer than the single mutant A (11,311), but when combined with H640L in the ADE triple mutant, site counts increased dramatically (>13,000). The ACDE quadruple mutant showed the largest shift, modifying more than 14,000 sites, a more than 2-fold increase compared to WT PAD4. Strikingly, over half of these sites overlapped with unique PAD2 targets, while almost none were exclusively shared with WT PAD4, indicating a drastic change in substrate preference.

**Figure 5.**
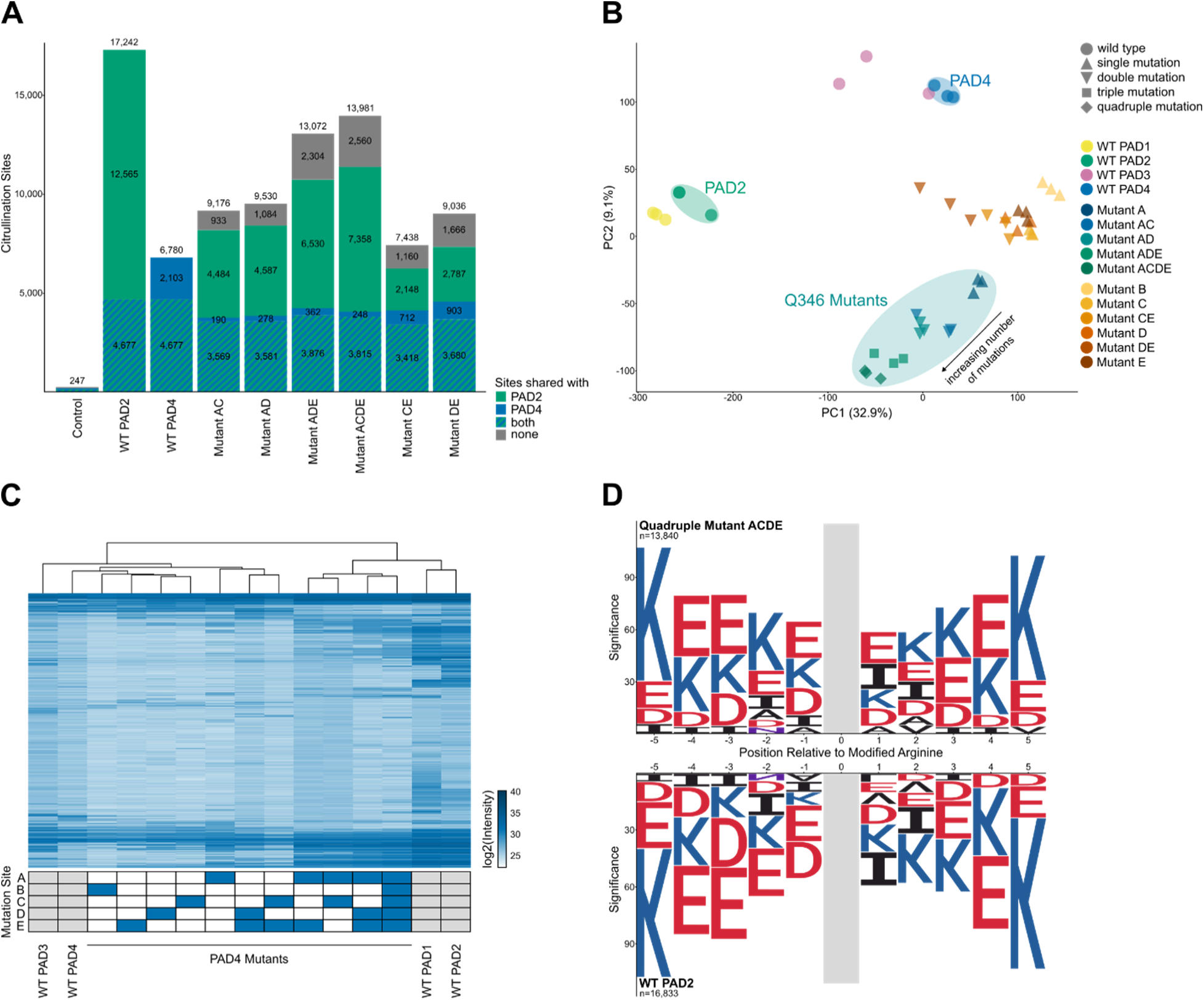
Multiple PAD4 mutations drive broad substrate recognition similar to PAD2. (A) Number of citrullination sites identified for each enzyme or PAD4 mutant compared to control (no enzyme). Shading indicates sites shared with wild-type PAD2, PAD4, or both. (B) Principal component analysis (PCA) of all citrullination sites identified across wild-type PAD1–4 and PAD4 mutants. (C) Heatmap of citrullination sites shared across all triplicates for wild-type PAD1–4 and PAD4 mutants. Rows and columns are hierarchically clustered by Euclidean distance. (D) Mirrored motif enrichment analyses comparing the PAD4 quadruple mutant (ACDE) with wild-type PAD2.

Global comparisons further confirmed this transition. Principal component analysis placed single and double mutants lacking Q346R (site A) close to WT PAD4 along PC1 (32.9% variance), with minor drift toward PAD2 in PC2 (9.1%). In contrast, all mutants including Q346R progressively approached PAD2 along both axes, with ACDE clustering closest to WT PAD2 (**Fig. 5B**). Hierarchical clustering of citrullination profiles mirrored this pattern (**Fig. 5C**): Site A-containing mutants (AC, AD, ADE, ACDE) grouped with PAD1 and PAD2, clearly separated from WT PAD4, PAD3, and the remaining variants.

Motif analysis revealed that combined mutations alter the specificity of PAD4. While double mutants retained elements of the canonical PAD4 motif, triple and quadruple mutants showed clear PAD2-like preferences (**Fig. 5D; Sup. Fig. 5**). In quadruple mutant ACDE, the dominant motif was essentially indistinguishable from PAD2, characterized by broader tolerance for acidic and basic flanking residues and diminished reliance on the strict –1 and +1 signature of PAD4 (**Fig. 5D**). Together, these results identify Q346, G403, R639, and H640 as central determinants of PAD4’s narrow specificity. Substituting these residues with PAD2 counterparts progressively broadens PAD4’s selectivity, expanding its repertoire and ultimately reprogramming it toward a PAD2-like substrate profile.

## DISCUSSION

In this study, we employed a highly sensitive mass spectrometry workflow combined with optimized data analysis pipelines to comprehensively and confidently characterize the substrate repertoires of human PAD1–4 isozymes. By identifying ∼30,000 citrullination sites from ∼5,500 proteins we provide, to the best of our knowledge, the most extensive mapping of PAD enzyme activity to date. This dataset represents a significant advance in understanding PAD substrate specificity, overcoming limitations of previous studies that relied on restricted sets of synthetic peptides^19,21,23^, a limited number of citrullination sites detected from unstimulated or unenriched biological samples^26,38,39^ and restricted or unclear isozyme responsibility for specific modification sites^20,22^.

Our proteome-wide analysis reveals new insights into the substrate selectivity landscape across the PAD family. While earlier studies primarily contrasted PAD2 and PAD4^23,24,40^, our inclusion of PAD1 and PAD3 expands the view to all catalytically active human isozymes. Notably, PAD1 and PAD2 cluster as a functional subgroup characterized by broader substrate tolerance, whereas PAD3 and PAD4 display greater selectivity. This division does not strictly align with known tissue distributions or physiological roles. Instead, it suggests that intrinsic biochemical features—rather than localization or biological function alone—drive substrate recognition preferences within the PAD family.

Mutagenesis experiments on PAD4 further revealed molecular determinants underlying isozyme-specific substrate recognition. Key substitutions (Q346R, G403S, R639F, H640L) near the active site were found to govern sequence motif preferences. For instance, replacement of polar or positively charged residues R639 and H640 (Mutants D and E) with hydrophobic counterparts from PAD2 (F and L, respectively) favored citrullination motifs enriched in hydrophobic residues (A, L, I, V). The most pronounced effect was observed with the Q346R substitution (Mutant A), where introduction of a positively charged arginine likely enhanced interactions with negatively charged sequence regions distal to the modified site. These findings are consistent with our motif analyses and structural interpretations. In contrast, mutation of R374, a residue previously implicated in substrate binding^36,37^, did not markedly alter motif preference, suggesting a role in substrate stabilization rather than in primary recognition.

A particularly striking outcome emerged when comparing enzymatic kinetics using the model substrate BAEE with proteome-wide citrullination activity. While BAEE-based activity assays accurately capture catalytic efficiency under defined conditions, they fail to reflect true substrate recognition in complex proteomes. For example, Mutant A (Q346R) exhibited comparable catalytic efficiency in BAEE assays but yielded the highest number of citrullination sites in the proteomic assay. Conversely, Mutant B (R374G) showed strong kinetic parameters but minimal gain in site count. This clear disconnect underscores the limitation of using simplified substrates to infer biological activity and highlights the need for proteome-scale validation when evaluating enzyme function or inhibitor design.

While our experimental design maximized sensitivity and reproducibility, several limitations warrant consideration. The *in vitro* lysate incubation system, although avoiding harsh detergents to preserve protein structures, remains an artificial environment. Extended incubation under saturating enzyme concentrations likely induced citrullination events not occurring physiologically. However, this trade-off was intentional, enabling an unbiased assessment of each enzyme’s intrinsic sequence preferences under standardized conditions, a critical prerequisite for comparative analysis. Other physiological factors, such as intracellular calcium gradients, isozyme localization, and interacting partners undoubtedly modulate PAD activity through allosteric or compartmental mechanisms but were beyond the scope of this study. Similarly, potential crosstalk with other post-translational modifications, including arginine methylation, was not addressed here, though prior studies indicate competition between these modifications^3,41–43^. Autocitrullination of PADs themselves may also regulate enzyme activity^44,45^. These regulatory layers will be important targets for future *in vivo* validation.

Beyond their mechanistic significance, our findings carry broader biological and translational implications. PAD activity is frequently dysregulated in disease contexts—including rheumatoid arthritis, cancer, and neurodegenerative disorders—where altered cellular environments (e.g., calcium flux, redox status, cofactor availability) may shift enzyme activity or substrate accessibility. Although such changes could influence which substrates are modified *in vivo*, they are unlikely to alter the intrinsic selectivity principles defined here. The distinct citrullination patterns observed across isozymes provide a framework to infer PAD activity from complex tissue datasets, enabling differentiation of PAD1/2-catalyzed from PAD3/4-catalyzed citrullinomes. Moreover, by identifying residues and structural features critical for substrate recognition, our study lays the groundwork for developing isozyme-selective inhibitors that target pathological citrullination without broadly suppressing all PAD activity.

In conclusion, this work delivers the most comprehensive mapping to date of substrate preferences across the active human PAD isozymes, identifies key molecular determinants of their selectivity, and reconciles biochemical and proteomic perspectives on enzyme specificity. Together, these findings redefine our understanding of PAD substrate recognition and provide a biochemical foundation for functional and therapeutic exploration of this important enzyme family.

## MATERIALS AND METHODS

### Cell Culture

HeLa cervical carcinoma CCL-2 (ATCC, Manassas, USA) and H4 neuroglioma (CLS Cell Lines Service, Eppelheim, Germany) cell lines were cultured in Dulbecco’s Modified Eagle Medium (DMEM) supplemented with 10% fetal bovine serum (FBS). Prior to collection, the cells were washed twice with ice-cold phosphate-buffered saline (PBS) to remove residual media. Lysis was performed using a gentle buffer containing 50 mM Tris-HCl (pH 7.5), 5% glycerol, 1.5 mM MgCl_2_, 150 mM NaCl, 1 mM Na_3_VO_4_, 25 mM NaF, 0.8% IGEPAL, and 1 mM dithiothreitol (DTT). The lysates were clarified by ultracentrifugation at 150,000 × g for 1 hour at 4 °C. Protein concentrations were determined using the Pierce BCA Protein Assay Kit (Thermo Fisher Scientific, Rockford, IL, USA) following the manufacturer’s recommended protocol.

### Purification of recombinant wild-type PAD2, PAD4, and PAD4 mutants

Human PAD2 and PAD4 cDNAs were cloned into pET19b and pQE30 vectors, respectively, to generate N-terminal His-tagged constructs. PAD4 mutants were generated by inverse PCR-based site-directed mutagenesis using the PAD4 construct as a template. Recombinant PAD2 was expressed in E. *coli* BL21(DE3)pLysS cells induced with 0.1 mM IPTG at OD_600_ 0.6, followed by incubation at 16°C, 150 rpm overnight. PAD4 and its mutants were expressed in E. *coli* JM109 cells induced with 1 mM IPTG at OD_600_ 0.8, and cultured at 25°C, 230 rpm overnight.

Cells were collected by centrifugation and disrupted by sonication in a buffer containing 30 mM Tris-HCl (pH 7.6), 500 mM NaCl, 2 mM β-mercaptoethanol, and 5 mM imidazole. The clarified lysates were incubated with Ni-NTA agarose resin (Sigma-Aldrich, St. Louis, MO, USA) for affinity purification of His-tagged proteins. After binding, the resin was sequentially washed with binding buffer (5 mM imidazole, 500 mM NaCl, 2 mM β-mercaptoethanol, 30 mM Tris-HCl, pH 7.6) followed by washing buffer (10 mM imidazole, 500 mM NaCl, 2 mM β-mercaptoethanol, 30 mM Tris-HCl, pH 7.6) to remove nonspecific contaminants. The target proteins were then eluted using elution buffer containing 250 mM imidazole, 500 mM NaCl, 2 mM β-mercaptoethanol, and 30 mM Tris-HCl (pH 7.6).

Eluted fractions were dialyzed against dialysis buffer (30 mM Tris-HCl, pH 7.6, 500 mM NaCl, 2 mM β-mercaptoethanol) and subsequently concentrated using Amicon Ultra-15 centrifugal filters (Millipore, Billerica, MA, USA; 50 kDa cutoff). The final protein preparations were mixed with 50% (v/v) glycerol, aliquoted, and stored at −80°C. Protein integrity and purity were assessed by SDS-PAGE analysis.

### PAD Activity Assay

PAD enzymatic activity was measured using a spectrophotometric assay coupled to glutamate dehydrogenase (GDH) under 25°C. The reaction mixture (200 μL per well, 96-well plate) contained Nα-benzoyl-L-arginine ethyl ester (BAEE) as substrate, 10 mM CaCl₂, 2.5 mM DTT, 8.5 mM α-ketoglutarate (α-KG), 0.22 mM NADH, and 8.4 U of GDH in 100 mM Tris-HCl (pH 7.6) and 150 mM NaCl. Reactions were performed at 25°C, and the decrease in NADH absorbance at 340 nm was monitored every 30 seconds using a Biotek Synergy HTX Plate Reader (Agilent Technologies, Santa Clara, CA, USA). Enzyme activity was defined as the amount of PAD catalyzing the oxidation of 1 μmol NADH per minute, with an extinction coefficient of 6.22 cm⁻¹ mM⁻¹ for NADH.

For kinetic analysis, initial reaction rates (*V*_0_) were determined at various concentrations of BAEE substrate ([*S*]). Absorbance changes at 340 nm were recorded in a 96-well plate, and reaction rates were corrected for the pathlength of the well (0.58 cm) using: *V*_0_=ΔA/Δt/0.58.

Kinetic parameters were obtained by fitting the corrected initial rates to the Michaelis–Menten equation using nonlinear regression in GraphPad Prism 7: *V*=*V*_max_*[*S*]/(*K*_m_+[*S*]). The turnover number (*k*_cat_) was calculated by dividing *V*_max_ by the molar concentration of PAD used in the assay: *k*_cat_ = *V*_max_/[*E*]_t_, where [*E*]_t_ represents the total enzyme concentration. And catalytic efficiency was expressed as *k*_cat_ / *K*_m_.

### *In vitro* Citrullination Assay

500 μg HeLa (conc.=3.9 μg/μL) (only wild-type experiments) or H4 (conc.=4.7 μg/μL) (wild-type, mutant, and time-dependent experiments) were diluted with 100 mM Tris (pH 7.6) to a final volume of 500 μL and supplemented with CaCl_2_ (10 mM final) and DTT (2.5 mM final). Recombinant PAD enzymes were prepared in 100 mM Tris and added to the reactions at concentrations adjusted to achieve identical enzymatic activity. Based on the activities determined by a spectrophotometric assay using BAEE as substrate^14^, the following amounts were applied to match the activity of PAD4 (produced in-house), which was added at a fixed enzyme-to-substrate protein weight ratio of 1:200: PAD1 (Cay10784, Biomol) at 1:79 enzyme:protein ratio, PAD3 (Cay10786, Biomol) at 1:119, and PAD2 (SAE0061, Sigma-Aldrich) at 2 units per reaction.

Analogously, in experiments with PAD4 mutants, enzymes were also added in ratios to match wild-type PAD4 activity: WT PAD4 at 1:150, Mutant A (Q346R) at 1:136, Mutant B (R374G) at 1:111, Mutant C (G403S) at 1:431, Mutant D (R639F) at 1:120, Mutant E (H640L) at 1:112, Mutant AC (Q346R|G403S) at 1:67, Mutant AD (Q346R|R639F) at 1:78, Mutant CE (G403S|H640L) at 1:417, Mutant DE (R639F|H640L) at 1:56, Mutant ADE (Q346R|R639F|H640L) at 1:56, Mutant ACDE (Q346R|G403S|R639F|H640L) at 1:65 enzyme:protein ratio.

For control samples, a matching volume of 100 mM Tris without enzyme was added (100 μL in wild-type, 5 μL in mutant experiments). Samples were incubated overnight at 37 °C with shaking at 600 rpm. For inactivation of PAD enzymes, the samples were subsequently heated for 10 min at 80°C. All enzyme reactions for wild-type and mutant experiments were performed in triplicates.

Time-course experiments were carried out in a similar fashion, using H4 lysate and in-house purified PAD2 and PAD4 isozymes at 1:45 And 1:200 enzyme:protein ratios, respectively. Samples were incubated for 10 min, 30 min, 1 h, 3 h or overnight at 37 °C with shaking at 600 rpm before heat-inactivation and freezing at -20°C until further processing. The control sample was prepared without any enzyme and directly frozen.

### Sample Preparation

To prepare samples for MS acquisition, they were reduced by adding DTT to a final concentration of 10 mM and incubating at 37°C for 45 min while shaking at 600 rpm. Alkylation was performed using CAA at 55 mM final and incubating at room temperature in the dark for 30 min. Trypsin was added at 1:50 enzyme:protein ratio and digestion carried out overnight at 37°C and 800 rpm shaking. Samples were then acidified with 1% formic acid and desalted using 50 mg Sep-Pak C18 SPE cartridges (Waters).

Basic reverse-phase fractionation was performed via centrifugation in self-constructed StageTips^46^ using a stack of six 8-gauge disks of C18 SPE material (Empore, CDS) and consecutively eluting peptide fractions with a 25 mM ammonium formate solution containing 5, 7.5, 10, 12.5, 15, 17.5 and 50% ACN. The flow-through and separated fractions were pooled to six fractions, and each fraction was again desalted using self-constructed StageTips (4 disks, 14-gauge) using the same C18 material. Samples were lyophilized and stored at -20°C until MS measurement.

### LC-MS/MS Data Acquisition

LC-MS/MS measurement was carried out on a Vanquish Neo LC system coupled to an Orbitrap Exploris 480 mass spectrometer (Thermo Fisher Scientific, Bremen, Germany). Samples were reconstituted in 12 μL of 0.1% FA, out of which 10 μL were injected with the LC loading buffer (0.1% FA) at a flow rate of 100 μL/min using the direct injection setup and fast loading mode. Separation was performed on an Acclaim PepMap RSLC C18 column (1 mm x 15 cm, C18, 2 μm, 100 Å, Thermo Fisher Scientific) temperature-controlled to 55 °C. Peptides were separated using a 60 min gradient of mobile phases A (0.1% FA, 3% DMSO in H_2_O) and B (0.1% FA, 3% DMSO in ACN), increasing linearly from 3% B to 24% B over the first 50 min, followed by another linear increase to 31% B between 50-60 min, at a flow rate of 50 μL/min. After completion of the gradient, the system was washed for 1.35 min at 90% B (100 μL/min) and equilibrated at 1 % B prior to the next run. Ionization in the heated ESI source was carried out at 3500 V, with sheath gas and auxiliary gas set to 32 and 5, respectively. The MS was operated in data-dependent acquisition (DDA) mode using the following parameters: MS1 scans were taken at a cycle time of 1.2 s with a resolution of 60,000, m/z range 360-1300 and RF lens 42%. MS2 scan filters included peptide profile for monoisotopic peak determination (MIPS), intensity threshold of 2.0e4, and charge state 2-4. Dynamic exclusion was enabled for 30s with a mass tolerance of 10 ppm. Precursors meeting the criteria were isolated within a window of 1.1 m/z, fragmented at an HCD collision energy of 28% and the MS2 scans measured with a resolution of 15,000, with the scan range defined by first mass set to 100 m/z. Fractions of the same sample were injected consecutively, with 15 min blanks acquired between samples, and instrument QCs measured after every set of triplicates.

### Database search and Prosit-Cit rescoring

The acquired raw data were searched with FragPipe (version 21.1, MSFragger version 4.0, IonQuant version 1.10.12, Philosopher version 5.1.0) against a fasta file of all human canonical sequences (downloaded from Uniprot 16.10.2024, 20,428 entries) supplemented with 20,428 decoy sequences using default peak matching settings. Strict trypsin was selected as digestion enzyme, allowing up to 3 missed cleavages, peptide length between 7-50 amino acids and a mass range of 500-5,000 Da. Carbamidomethylation of cysteine was set as fixed modification, oxidation of methionine and acetylation of protein N-term as variable modifications. Additionally, up to 3 occurrences of citrullination on arginine (+0.9840 Da), and deamidation on asparagine or glutamine (+0.9840 Da) were allowed, with a total maximum of 3 variable modifications per peptide. Rescoring using MSBooster was disabled. For full proteome data, FDR cutoffs were set to 0.01 at peptide and protein level, and sequential algorithm enabled. For citrullination data, protein interference minimal probability was set to 0, and FDR filters were set to 1 at ion, PSM, peptide and protein level. All other search parameters were kept at default values. To increase confidence in citrullination identifications, the data were rescored using the Prosit-Cit algorithm within Oktoberfest^32^ and filtered for a q-value of 0.01 at PSM level for FDR control. Resulting PSMs were mapped back to the intensity values provided in the FragPipe output.

### Data preprocessing

From the PSM data frame, any modified sequences with C-terminal citrullination were removed. Sequences detected in multiple fractions or charge states were combined by adding up their intensity values. For sequences with multiple site annotations, the intensity values were duplicated for each possible site assignment, and identical sites combined by summing up intensities to finally obtain a single intensity value per site. One raw file of one of the HeLa PAD2 samples was corrupted, so that this replicate had to be excluded from data analysis, leaving only two replicates for this condition.

For most downstream data analysis of endpoint wild-type and mutant datasets (except principal component analysis and heatmaps), only citrullination sites were considered that were routinely detected in all replicates of the respective condition, with the mean intensity value across replicates being used. For principal component analysis and heatmaps, a Perseus style imputation was performed, replacing missing values by random numbers drawn from a normal distribution of 1.8 standard deviation down shift and a width of 0.3 for each sample. For heatmaps on time-dependent experiments, only sites identified in either 3 h or 16 h timepoints were included, and the filtered datasets imputed as described above. After clustering citrullination sites (Euclidean distance), only those clusters showing early, intermediate or late modification were visualized.

### Data analysis and visualization

#### Analysis in R

Custom scripts were used for further data analysis and visualization in RStudio, making use of packages ggplot2 (v 3.4.2) for barplots, violinplots and boxplots, pheatmap (v 1.0.12) for heatmaps and clustering, ggridges (0.5.6) for ridgeplots, and eulerr (v 7.0.1) for Venn diagrams. R Stats package (v 4.2.0) was used for clustering and PCA analyses.

#### Motif analysis

Motif analysis was performed by submitting identified citrullination sites (sequences including 5 amino acids to either side of the modified arginine) to the pLogo online probability logo generator^33^. Fasta files of all identified proteins in the respective cell lines (HeLa or H4) were uploaded as background proteome. Only those amino acids significantly over-represented in the sample compared to the respective background (p = 0.05, Bonferroni corrected) as identified by pLogo were then visualized using the ggseqlogo R package (v 0.2)^47^.

#### Structure prediction

Secondary structures of identified citrullinated arginines were predicted using NetSurfP-3.0 web server (DTU Health Tech, Denmark)^34^. Canonical fasta sequences of all proteins with a citrullination site found in the H4 or HeLa dataset were retrieved from Uniprot using the ID mapping tool to human UniProtKB/Swiss-Prot database (downloaded 10.04.2025). Protein sequences with more than 5,000 amino acids, or containing selenocysteine or pyrrolysine were removed. Remaining sequences were submitted to the NetSurfP-3.0 online tool in batches, and the output downloaded in csv format. The prediction values for arginine residues was mapped to the corresponding citrullination sites. For secondary structure and relative solvent accessibility comparisons, all arginines from proteins identified with 1%FDR from the H4 and HeLa data sets served as background.

AlphaFold predictions of the entire human proteome were downloaded from AlphaFold Protein Structure Database^35^ (v 4, downloaded 10.04.2025, containing 23,391 predicted structures). Proteins larger than 2,700 amino acids were removed, since they are split up for structure prediction and contain multiple prediction values for the same site. pLDDT values for identified citrullination sites were extracted and compared to the background of all predicted values of arginines within proteins identified with 1% FDR from the H4 and HeLa data sets.

## DATA AVAILABILITY

The mass spectrometry proteomics data have been deposited to the ProteomeXchange Consortium via the PRIDE^48^ partner repository with the dataset identifier PXD071247.

## ACKNOWLEDGEMENTS

The authors would like to express their gratitude to all members of the Lee Lab, the Bavarian Center for Biomolecular Mass Spectrometry (BayBioMS), and the Chair of Proteomics and Bioanalytics for their valuable assistance and insightful discussions. This work was funded by the Federal Ministry of Education and Research (FKZ031L0215; YIG-SysNS), the Ministry of Science and Technology, Taiwan (NSTC 111-2311-B-005 -003) and an ERC Starting Grant (101077037; ORIGIN). The Exploris 480 mass spectrometer was funded in part by the German Research Foundation (DFG-INST 36/171-1 FUGG). The graphical abstract and illustrations were created with BioRender.com.

## AUTHOR CONTRIBUTIONS

S.L. and C.-Y.L. designed the research. S.L., Y.-F.Y., K.-H.C., and R.M.G. performed the experiments and collected the data. W.G. and S.L. processed the raw data. S.L. and C.-Y.L. carried out the formal analysis and interpreted the results. M.W. and H.-C.H. provided analysis tools, research materials, and instrumentation. C.-Y.L. supervised the project, secured funding, and revised the manuscript. S.L. drafted the manuscript, and all authors contributed to reviewing and editing the final text and approved the final version.

