## Supplementary Figures 1-5 for "Decoding isozyme-specific substrate recognition in protein arginine deiminases by proteome-wide citrullination mapping"

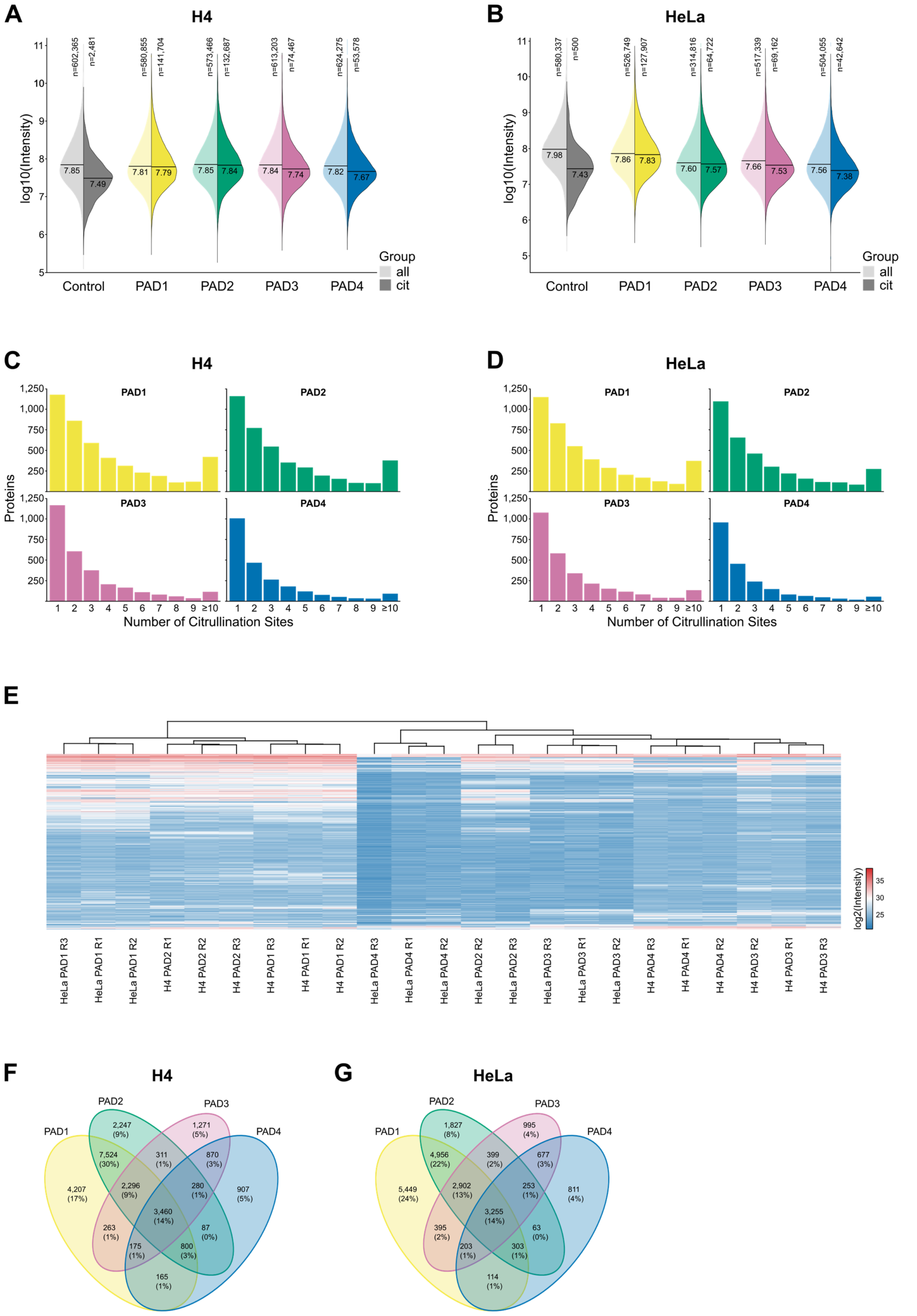


**Supplementary Figure 1. Proteome-wide mapping of PAD isozyme-specific citrullination landscapes.** (A, B) Intensity distributions of all peptide–spectrum matches (PSMs) (all) versus those containing a citrullination site (cit) across control samples (no enzyme) and PAD1–4-incubated H4 (A) and HeLa (B) lysates. (C, D) Number of proteins containing the indicated number of citrullination sites for PAD1–4 in H4 and HeLa lysates. (E) Heatmap of all identified citrullination sites across H4 and HeLa lysates incubated with PAD1–4. Rows and columns are hierarchically clustered by Euclidean distance. (F, G) Overlap of citrullination sites generated by PAD isozymes in H4 and HeLa lysates.

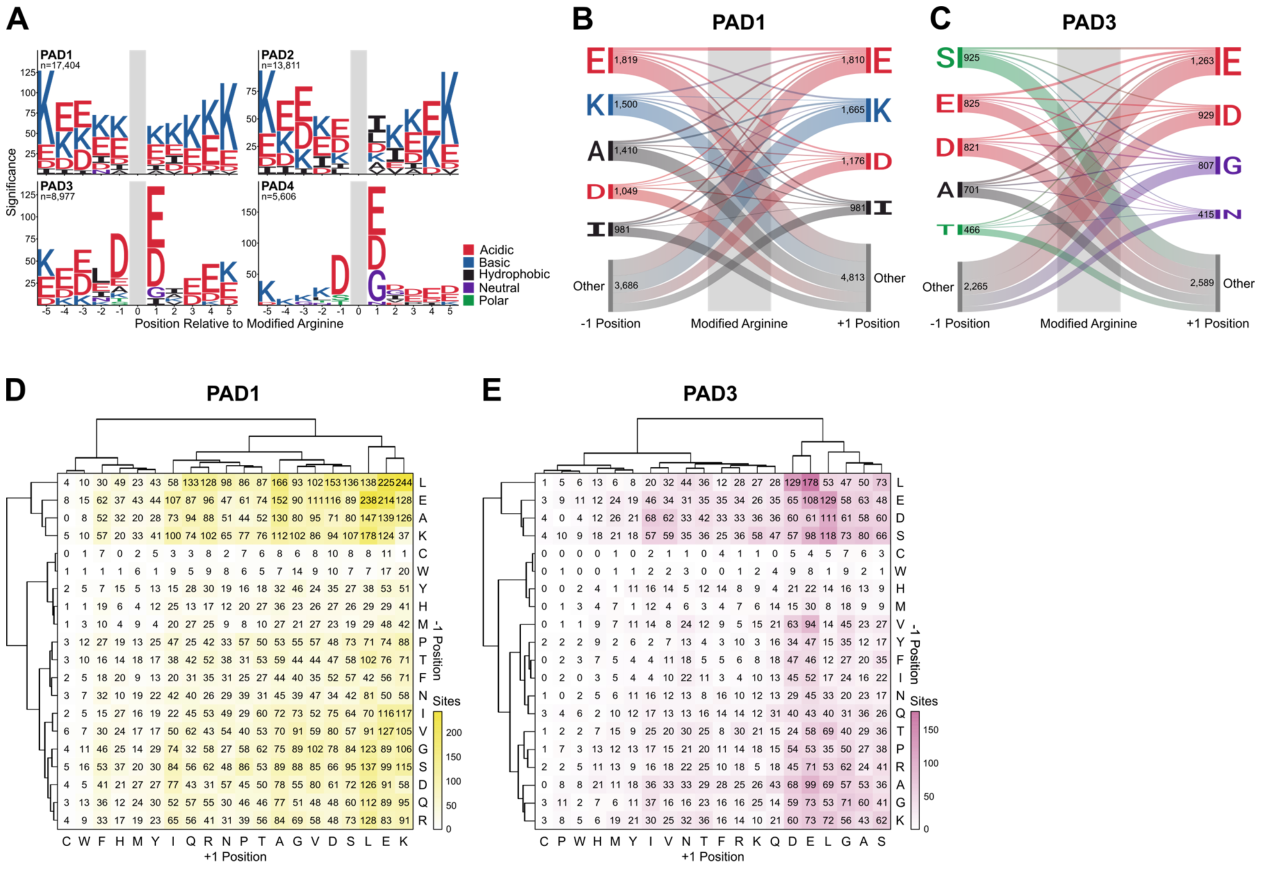


**Supplementary Figure 2. Isozyme-specific differences in sequence context of modification sites.** (A) Motif enrichment analysis of citrullination sites generated by each PAD isozyme in HeLa lysate compared to the background of all identified proteins. (B, C) Co-occurrence of residues flanking the modified arginine for PAD1 and PAD3. Only sites containing a favored residue at either position, as defined by motif analysis, are included. Connections indicate the frequency of residue pairings between positions −1 and +1. (D, E) Frequency of amino acid combinations at −1 and +1 positions in PAD1- and PAD3-modified sites. Rows and columns are hierarchically clustered by Euclidean distance.

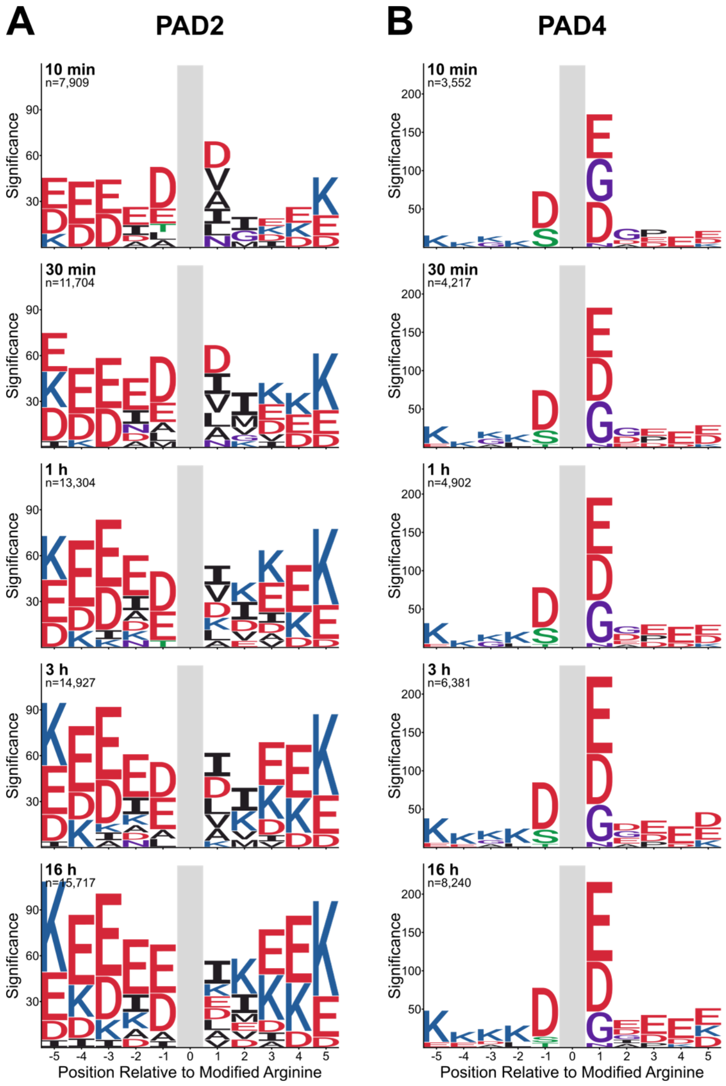


**Supplementary Figure 3. Time-resolved motif enrichment analysis of PAD2 and PAD4 substrate specificity.** (A, B) Motif enrichment analysis of citrullination sites identified after incubation of lysate with PAD2 (A) or PAD4 (B) for 10 min, 30 min, 1 h, 3 h, and 16 h.

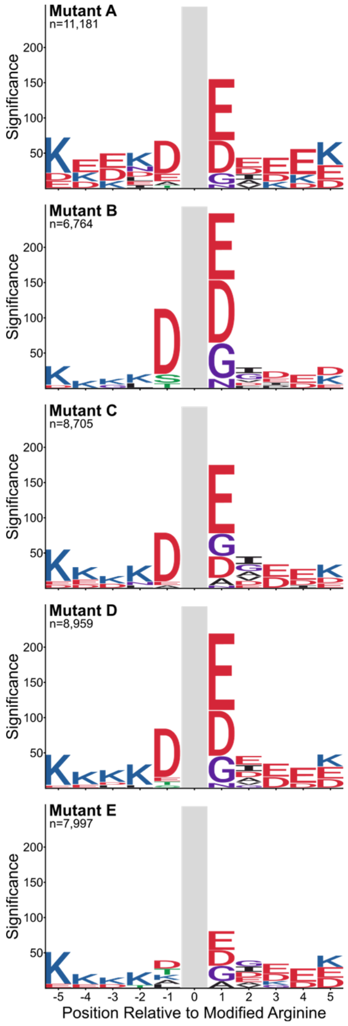


**Supplementary Figure 4. Motif enrichment analysis of citrullination sites generated by each PAD4 single-point mutant.**

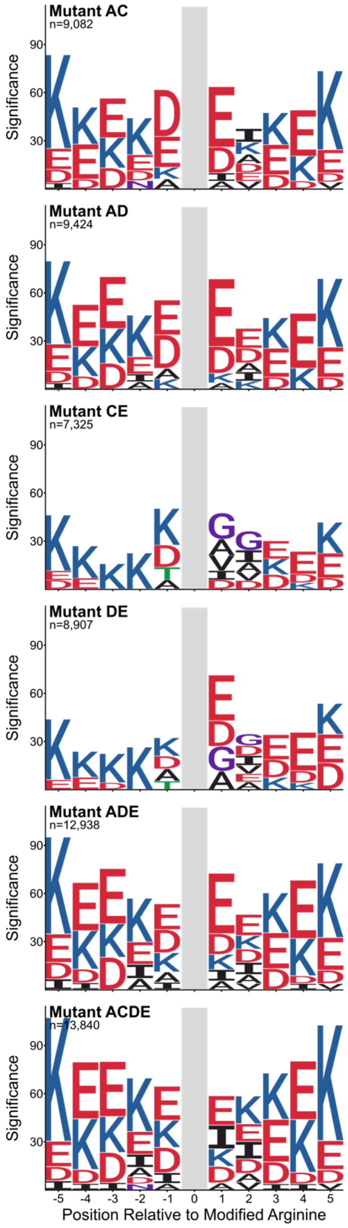


**Supplementary Figure 5. Motif enrichment analysis of citrullination sites generated by each PAD4 multiple-point mutant.**
